# Relevance of host cell surface glycan structure for cell specificity of influenza A virus

**DOI:** 10.1101/203349

**Authors:** Markus Kastner, Andreas Karner, Rong Zhu, Qiang Huang, Dandan Zhang, Jianping Liu, Andreas Geissner, Anne Sadewasser, Markus Lesch, Xenia Wörmann, Alexander Karlas, Peter Seeberger, Thorsten Wolff, Peter Hinterdorfer, Andreas Herrmann, Christian Sieben

## Abstract

Influenza A viruses (IAV) initiate infection *via* binding of the viral hemagglutinin (HA) to sialylated glycan receptors on host cells. HAs receptor specificity towards sialic acid (SA) is well studied and clearly critical for virus infection, but the contribution of the highly complex cellular plasma membrane to the cellular specificity remains elusive. In addition, some studies indicated that other host cell factors such as the epidermal growth factor receptor might contribute to the initial virus-cell contact and further downstream signaling^1^.

Here we use two complementary methods, glycan arrays and single-virus force spectroscopy (SVFS) to compare influenza virus receptor specificity with actual host cell binding. Unexpectedly, our study reveals that HAs receptor binding preference does not necessarily reflect virus-cell specificity. We propose SVFS as a tool to elucidate the cell binding preference of IAV thereby including the complex environment of sialylated receptors within the plasma membrane of living cells.

## 1. Introduction

Influenza A viruses circulate in aquatic birds, their large natural host reservoir, but have also established stable lineages in various mammalian species such as pigs. Although animal influenza viruses are usually confined to their natural host species, they can cause zoonotic infections in humans on rare occasions^2^. Such trans-species transmissions can result in clinically severe or even fatal respiratory disease in humans as illustrated by the outbreaks of avian-origin H7N9 subtype viruses in China occurring since 2013^3^. Zoonotic transmission events can, in fact, largely influence the epidemiology of human influenza directly if the virus succeeds to spread among humans as was observed in 2009 for the pandemic swine-origin H1N1 strain (pdmH1N1)^4^. Although the genetic requirements for crossing the species barrier are still incompletely understood, it is accepted that interspecies transmission of influenza A viruses partially depends on the capability of viral hemagglutinin (HA) to recognize specific sialylated-glycan receptors on the host cell surface. In general, HA of avian viruses preferentially binds to α-2,3-linked sialic acid (SA) (avian-type receptor) ^5^ whereas HA of human-adapted strains strongly bind to terminal α-2,6-linked SA (human-type receptor) ^5^. Several studies have determined that alterations in HA receptor binding specificity are a crucial step in host adaptation and interspecies transmission for several IAV subtypes ^6, 7, 8^. However, it is not well established if those adaptive mutations (1) provide an actual advantage in virus-cell binding during entry, or (2) whether they are necessary to confer transmission (i.e. by evading decoy receptors lining the human airway mucus) or (3) to avoid triggering of innate immune signaling. Regarding the first point, several studies suggest a much higher complexity of virus–cell interaction beyond the level of HA–SA binding (for a review see^9^). Consequently, it was hypothesized that human influenza viruses bind to a more structurally diverse set of SA linked carbohydrates than avian viruses which goes beyond the general preference of α-2,3 or α-2,6 linkage.

Currently, glycan arrays with libraries of synthesized glycan structures are widely utilized for the characterization of IAV glycan specificity. In particular, due to direct exposure of receptors on the array, sialic acid specificity can be studied with high precision on structural glycan properties. However, the cellular glycome has been recently studied for human and swine respiratory tract tissue, showing that its complexity might not be well represented by current glycan arrays ^13, 14^. Indeed, influenza virus infection in the absence of sialic acid suggests other possible attachment factors involved in virus binding. Candidates molecules are C-type lectins (L-SIGN and DC-SIGN), which were found to participate in influenza virus attachment independent of SA specificity ^12^. Hence, complementary approaches to directly assess viral receptor specificity within the complex environment of the cell surface are necessary to reach a more comprehensive understanding of the initial stage of virus infection. As we have recently shown, atomic force microscopy (AFM)-based single-virus force spectroscopy (SVFS) allows to measure the binding of individual IAV to living host cells at the molecular level ^15, 16, 17^. In this type of analysis, intact influenza viruses are covalently attached to AFM cantilevers, which are then lowered on single living cells (Fig. 1). Cycles between cantilever-cell approach, cell binding and cantilever retraction, allow direct characterization of virus cell binding, while revealing kinetic and thermodynamic properties of the interactions ^15, 16^. Thus, SVFS allows to investigate virus-cell binding in an experimental system that closely mimics the natural situation.

**Figure 1.**
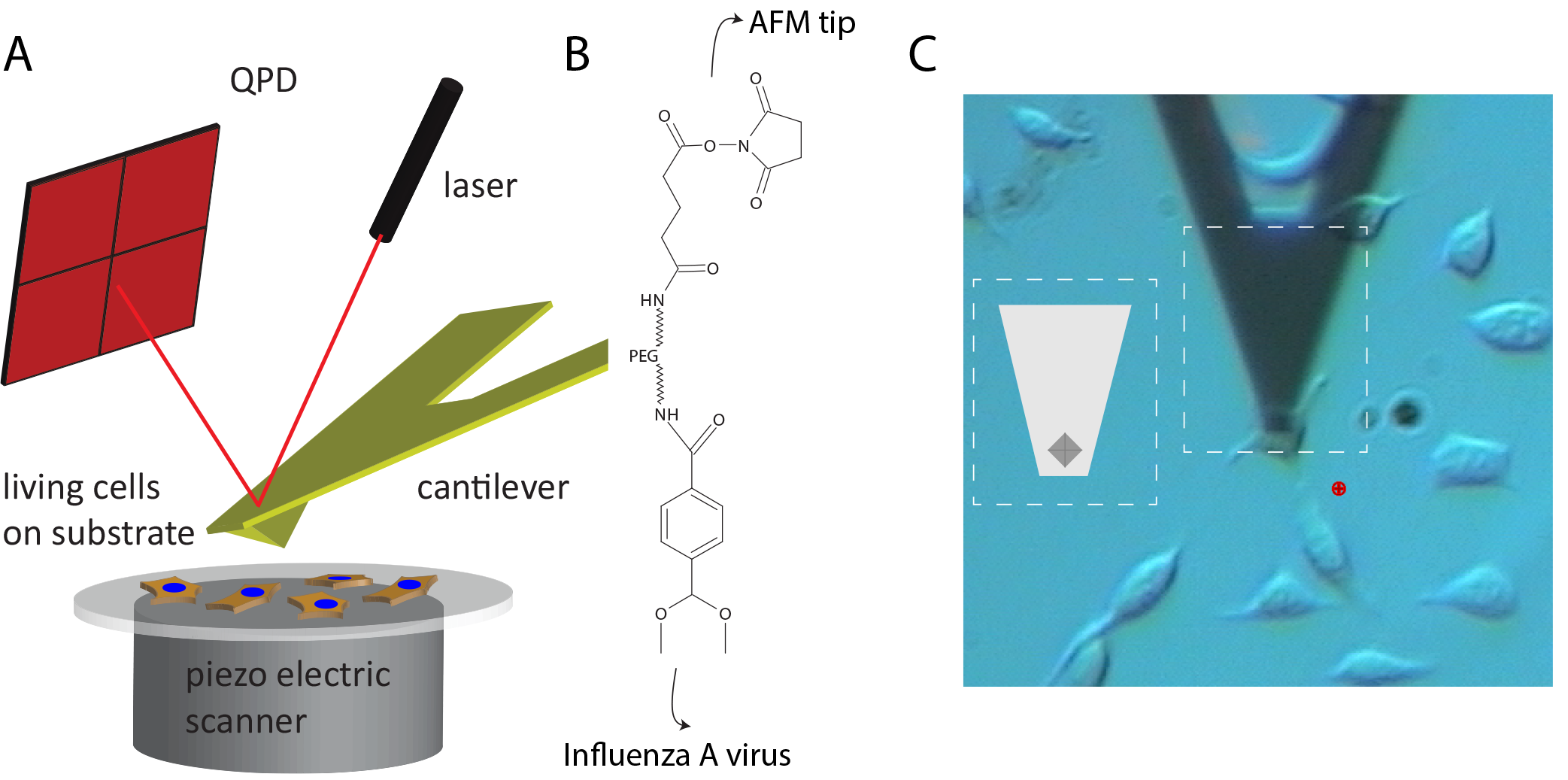
Schematic diagram of the SVFS experimental setup using atomic force microscopy (AFM). **(A)** General principle of AFM-based SVFS. Cells grow in a plastic culture dish that is attached to three step motors that allow movement with high accuracy. The cantilever acts as a Hookean spring and hence bending can be translated into applied force. The force-induced deflection of the cantilever is measured by pointing a laser on the back of the cantilever while detecting the reflection on a quadrant photo diode (QPD). **(B)** For SVFS, influenza A virions are covalently attached to the cantilever using an acetal-PEG_800_-NHS crosslinker ^39^. (**C**) The cantilever is lowered on a single cell until touching the cell surface. The combination with light microscopy, allows identification of the cantilever with its pyramidal cantilever tip (**C**, inset shows a graphical illustration) and thereby precise positioning. Subsequently, the cantilever is retracted at a defined velocity v. In case of an interaction, the cantilever will bend towards the sample until the underlying bond fails and the cantilever returns into the zero-force position (see also Fig. 2a).

Here, by using a set of five different influenza A virus strains, we systematically address whether virus-cell specific binding patterns as determined by SVFS are reflected in their receptor specificity observed by glycan array analysis. Our data indicate that results obtained from *in vitro* glycan arrays may not be directly transferred to virus-cell binding. We suggest that host cell specificity does not solely depend on the sialic acid configuration of the cell surface, but is more complex and depends on the specific environment of the receptor and possibly involves additional attachment factors or co-receptors with yet unknown functional roles.

## 2. Results

### Receptor specificity of influenza A virus strains studied by glycan arrays

To investigate and compare the SA receptor specificity of different virus strains, we performed an *in vitro* glycan array study utilizing a library of 15 glycans (Fig. 2). Regarding specific IAV receptors, our library included three α-2,3-linked (avian-type) SA conjugates as well as three α-2,6-linked (human-type) SA conjugates. The glycan number nine, sialyl-Lewis^X^ (SLe^X^) has due to its fucosylation a different topology and was, although α-2,3-linked to SA, separated from the two groups shown on the left side in Fig. 2.

We found that the zoonotic AH1 strain and the pandemic H1N1 virus recognized all six SA conjugates with a preference for α-2,6-linked (human-type) receptors. Such a dual-binding behavior was already observed before for pdmH1N1 (A/California/04/2009 and A/Hamburg/5/2009). The receptor binding preference of the human H7N9 isolate A/Anhui/1/2013 (AH1) is still under debate. It was shown that AH1 exhibits increased human receptor binding while still preferring avian receptors ^20^, others reported on the specificity for human-type receptors ^21^. Interestingly, when we looked at the cumulative difference between the tested virus strains (Fig. S1), receptor specificity of AH1 and pdmH1N1 was most similar (i.e. the lowest difference).

**Figure 2.**
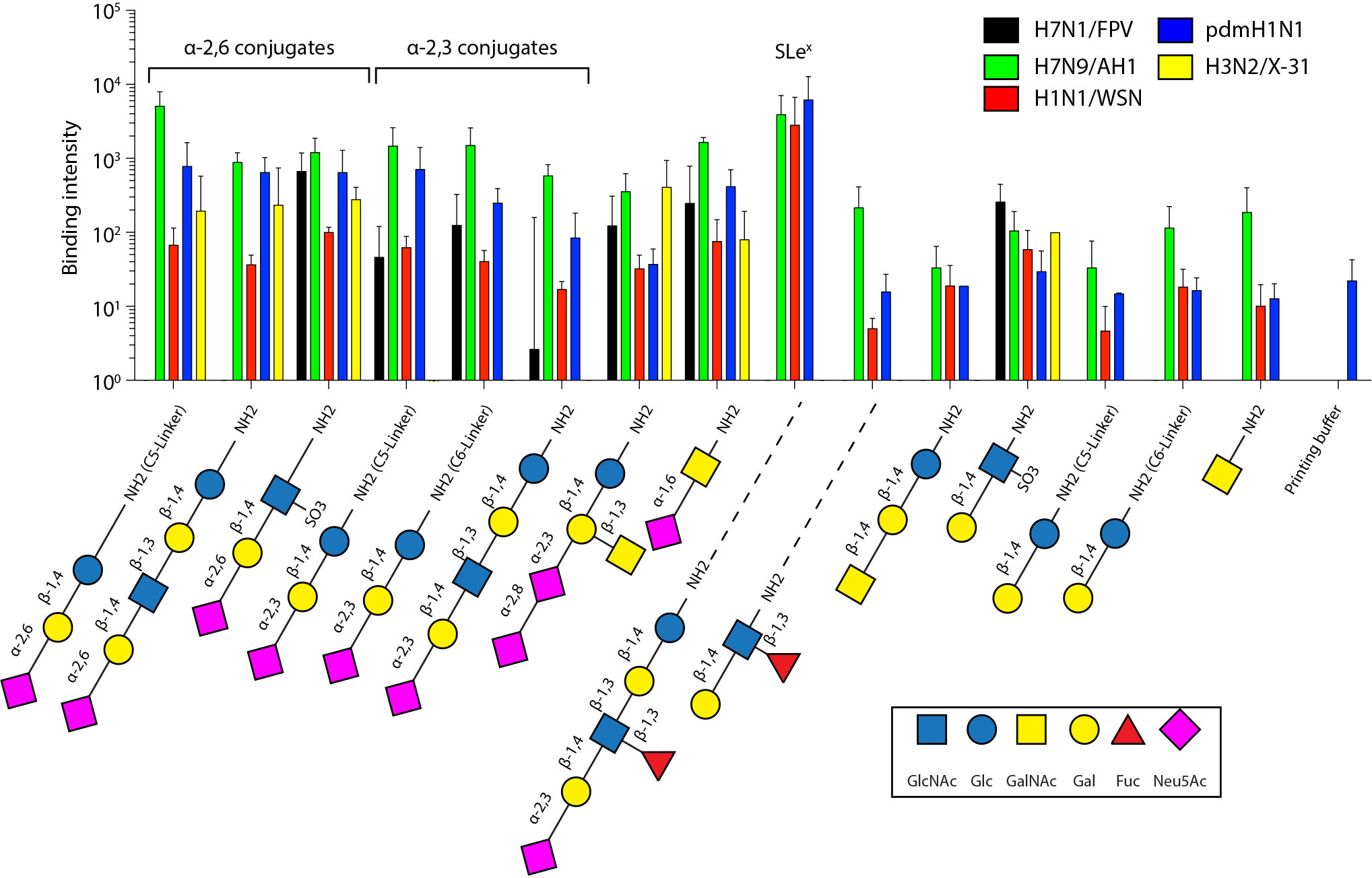
Binding characteristics of the indicated viruses to sialic acid-conjugated receptors quantified by glycan arrays. Equal amounts of the indicated viruses were bound to glycan arrays, spotted with 15 different sialic acids and printing buffer as negative control. Staining of bound viruses was achieved using a NP-specific primary antibody and a Cy3-coupled secondary antibody. The results represent the mean + SD for two independent experiments.

FPV recognized all three avian-type conjugates, but bound only one human-type conjugate, which is in line with previous findings using glycan arrays ^22^.

We further tested H3N2/X31 as well as H1N1/WSN. H3N2/X31 carries the HA of the human pathogenic strain A/Aichi/68, which was also previously shown to prefer α-2,6-linked (human-type) receptors ^23^. In line with that, we found that H3N2/X31 only recognized human-type SA conjugates on the glycan array. The lab-adapted H1N1/WSN was previously shown to prefer α-2,6-linked (human-type) receptors over α-2,3-linked SA on re-sialylated erythrocytes ^24^. In our hands H1N1/WSN bound all six receptors with no obvious preference. Its worth mentioning that among all glycans, we observed the strongest binding for pdmH1N1, H1N1/WSN and AH1 to the kinked, fucosylated glycan SLe^X^ (Fig. 1). SLe^X^ is well-known as an integrin receptor on leucocytes ^25^, but was also shown to be recognized by different IAV subtypes ^26^. However, FPV was shown to bind only the sulfated form of SLe^X^, which is in line with our findings ^22^.

### Cell specificity of influenza A virus strains studied by SVFS

Next, we used SVFS to characterize virus binding to two different cell types: First, we studied living A549 cells, a model cell line derived from the lower human respiratory tract, expressing both major SA receptor types on the cell surface as shown by lectin binding ^15, 18^. Secondly, CHO cells lack an α-2,6-specific sialyltransferase and only express α-2,3-linked SA. Hence, we chose them as a comparative model for studying viral binding to cells displaying only avian-type receptors ^15^. For SVFS, intact viruses were covalently attached to AFM cantilevers as previously reported (Fig. 1)^17^. Binding to cells was measured in a dynamic range of increasing loading rates, i.e. pulling velocities to determine the dissociation rate at zero force *k*_*off*_. Unbinding events were recorded and analyzed to obtain the rupture force *F* as well as the effective spring constant *k*_*eff*_, defined as the slope of the force-distance curve at rupture (Fig. 3a). From *k*_*eff*_, the loading rate *r* (force per time) was calculated by multiplication with the retraction velocity *ν*. Notably, we used an adapted data analysis procedure, which takes the variable local conditions of a living cell surface into account ^27^. Briefly, although the loading rate *r* should be constant for a given pulling speed *ν*, recent studies have shown that the heterogeneity of a living cell surface leads to a broad distribution of observed loading rates ^27^. Hence, to account for this effect, our approach does not rely on binning of loading rates, but takes each individual force-distance curve into account (Fig. 3c). By fitting the force spectra to a single energy barrier model (Fig. 3c, d, see *Materials and Methods*), we obtained the thermodynamic properties of the interaction such as the dissociation rate *k*_*off*_, and the separation of the receptor-bound state to the energy barrier *x*_*u*_ (summarized in table 1). The dissociation rate *k*_*off*_ and its reciprocal, the bond lifetime *τ*_*off*_ provide information about the stability of the underlying virus-cell interaction. The results for all virus-cell interaction pairs are illustrated in Fig. 2d. For details, please see *Materials and Methods*.

**Table 1.** Dissociation rate *k*_*off*_, separation from the energy barrier *x*_u_, and average bond lifetime *τ*_off_ obtained by fitting the SVFS data to a single energy barrier binding model as described in Methods (see also Fig. 2D).

| Cell (receptor type) | $x_u$ (Å) | $k_{\text{off}}$ (s<sup>-1</sup>) | $\tau_{\text{off}}$ (s) |
| --- | --- | --- | --- |
| **Virus AH1 (H7N9)** |  |  |  |
| CHO (*avian-like*) | $13.2 \pm 0.016$ | $1.17 \pm 0.001$ | 0.85 |
| A549 (*human-like*) | $9.24 \pm 0.006$ | $0.69 \pm 0.0008$ | 1.43 |
| **Virus FPV (H7N1)** |  |  |  |
| CHO (*avian-like*) | $2.40 \pm 0.004$ | $0.33 \pm 0.001$ | 3.03 |
| A549 (*human-like*) | $5.74 \pm 0.016$ | $0.24 \pm 0.001$ | 4.15 |
| **Virus pdmH1N1** |  |  |  |
| CHO (*avian-like*) | $12.5 \pm 0.004$ | $0.19 \pm 0.001$ | 5.20 |
| A549 (*human-like*) | $5.2 \pm 0.016$ | $0.2 \pm 0.001$ | 5.00 |
| **Virus X31 (H3N2)** |  |  |  |
| CHO (*avian-like*) | $9.54 \pm 0.18$ | $0.66 \pm 0.05$ | 1.51 |
| A549 (*human-like*) | $6.42 \pm 0.09$ | $1.27 \pm 0.07$ | 0.78 |
| **Virus WSN (H1N1)** |  |  |  |
| CHO (*avian-like*) | $2.77 \pm 0.04$ | $0.62 \pm 0.04$ | 1.61 |
| A549 (*human-like*) | $2.67 \pm 0.04$ | $0.85 \pm 0.05$ | 1.18 |

**Figure 3.**
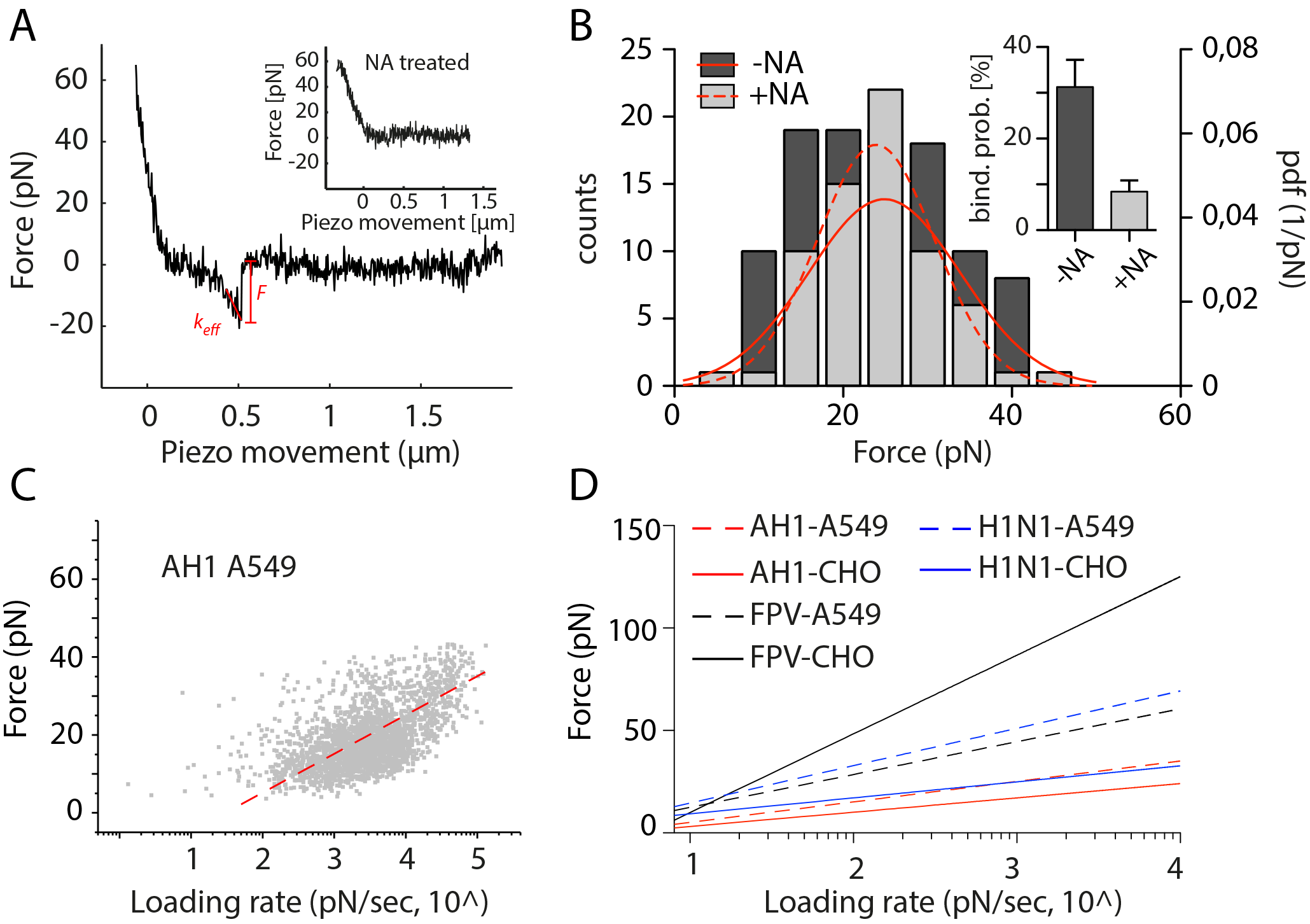
SVFS measurements of H7 and pdmH1N1 viruses interacting with receptors on living cells. (**A**) Force trace of H7N9 AH1 virus-cell interactions measured by AFM-based SVFS showing a characteristic single unbinding event. After treating the cells with neuraminidase (NA), the binding probability was strongly decreased (see also inset in B), causing a high number of force traces showing no interaction (inset in a). (**B**) Force histogram (left Y axis) and overlaid force probability density function (pdf, right Y axis) of AH1 virus-A549 cell interaction before and after NA treatment. The observed force values were found to be very similar, but the binding probability was strongly decreased (inset). (**C**) Scatter plots showing unbinding force *F* plotted against the loading rate *r* of every individual force curve from AH1 virus-A549 cell interaction. The red dotted line shows the fitting to the single energy barrier model (see Methods). (**D**) Overview of the fittings used to determine values for *k*_*off*_ and *x*_u_ (see table 1 and Methods) for all virus-cell combinations.

Fig. 3b shows a typical rupture force histogram and the accompanying probability density function (pdf) of the interaction between influenza AH1 and A549 cells with a binding probability of 29.7 % (*ν* = 500 nm/sec). The pdf shows a single peak at ~23 pN indicating specific interaction (red curve in Fig. 3b). After cell surface SA deprivation by neuraminidase (NA) treatment, the binding probability was reduced to 5-13 % while the pdf peak position was unchanged (red curves and inset in Fig. 3b). This verifies the specificity of our measurement for receptor interaction. The binned histograms are shown for comparison along with the fitted pdf.

For AH1, we observed pronounced binding to both tested cell lines, with rupture forces, between 10 and 100 pN depending on the applied loading rate (Fig. 3c, d). However, we found an about 40 % reduced dissociation rate for A549 compared to CHO cells, indicating preferential binding of human-type cell surfaces. We confirm this binding preference of AH1 by measuring binding to living MDCK cells, which express, similar to A549 cells, both human-and avian-type receptors. We observed preferential binding to MDCK cells compared to CHO cells (Fig. S2). For FPV, we observed about three times lower dissociation rates compared to AH1, with preferential binding to A549 (see table 1). pdmH1N1 virus showed similar dissociation rates as FPV, but without pronounced cell type preference. H3N2/X31 as well as H1N1/WSN were already studied by SVFS in our previous study^15^ but reanalyzed using the improved fitting procedure described above. The fitting values are reported in table 1. H3N2/X31 showed stronger attachment to CHO cells, while binding of H1N1/WSN to A549 and CHO cells was almost identical ^15^.

## 3. Discussion

Among other methods, solid-phase binding assays or glycan arrays represent a widely used state-of-the-art way to analyze HA receptor specificity ^28^. The desired ligand is coupled to a flat surface and can either be probed with intact viruses ^7, 8^ or purified HA ^19^, which is then detected using antibody binding. This makes them a powerful tool to screen large glycan subsets. Choosing the right library is critical and since recent glycomics studies indicate a large heterogeneity of host cell-specific glycans, this choice is not easily made ^14, Byrd-Leotis, 2014 #5^. Also, the presentation (i.e. orientation and density) of the glycan is critical ^29^, a factor that can be estimated by performing avidity studies along with the glycan array ^30^. In our study, we have performed glycan array binding to evaluate the receptor specificity of various virus strains along with testing their cell specificity using SVFS. While our glycan array results are largely in line with previous findings, SVFS results, as summarized in table 1 and Fig. 4a, suggest that HA’s preference for human or avian-type receptors does not necessarily correlate with expected binding patterns to cell lines modelling the surfaces of human or avian cells (see below). We compared the specificity of virus binding measured in a glycan array (i.e. receptor specificity) with that measured by SVFS (i.e. cell specificity). For pdmH1N1 and H1N1/WSN, we found that the SVFS data (for H1N1/WSN see ^15^) are in good agreement with results obtained from glycan array binding as neither strain displayed strong preference for human or avian-type receptors or a particular cell model.

**Figure 4.**
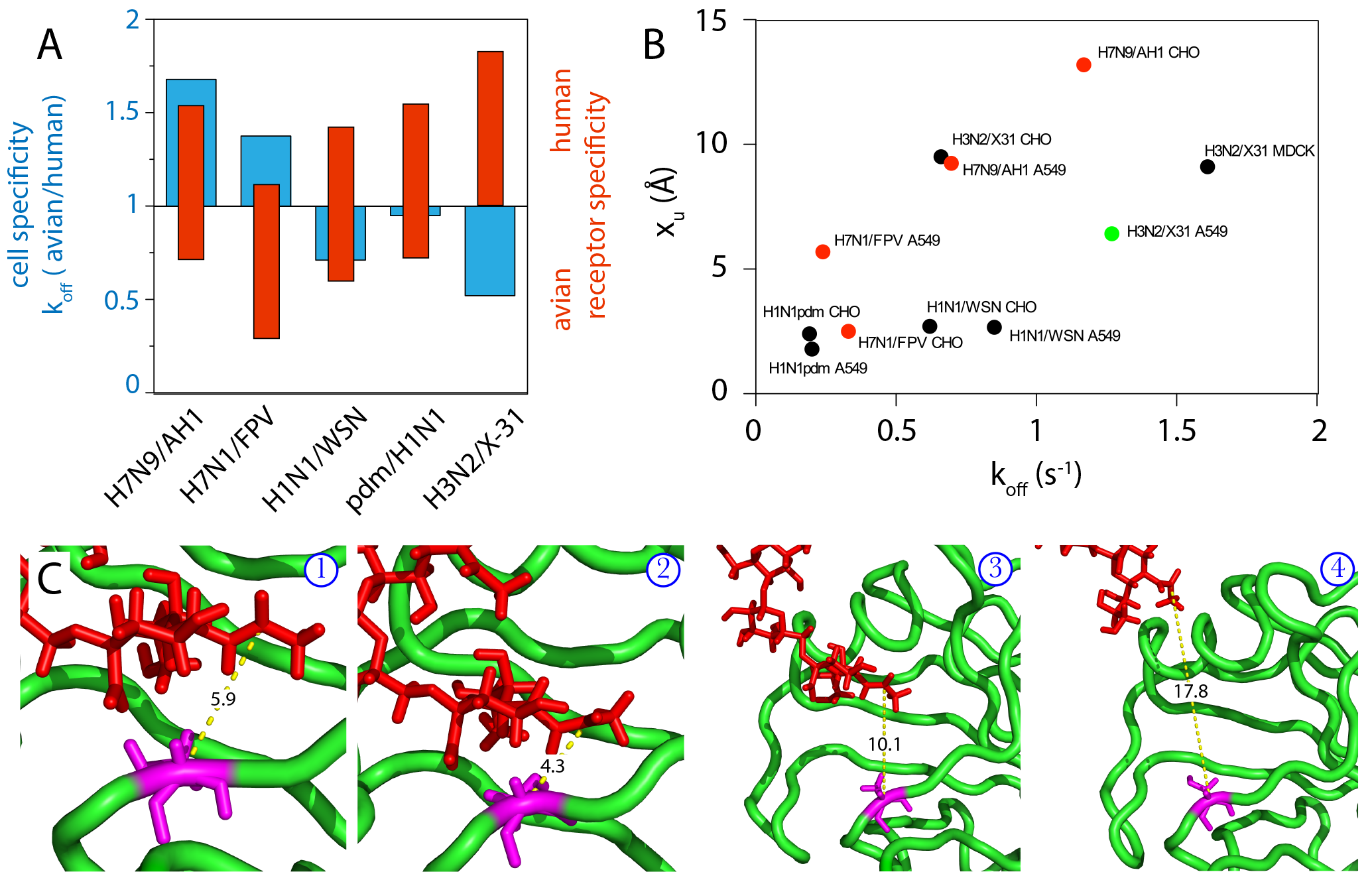
Comparing cell with receptor specificity and summary of the thermodynamic parameters obtained from SVFS. (**A**) To allow comparison between cell and receptor specificity, we show results from SVFS (blue bars) as Δk_off_ (avian/human) as well as results obtained from glycan arrays (red bars). The red bars indicating receptor specificity were placed according to the number of recognized glycans as well as the strength of binding, as discussed in section 2. (**B**) Correlation of *k*_*off*_ and *x*_*u*_ shows no apparent clustering for H7 (red) or H1 (black) viruses indicating a very dynamic interaction (as shown in ^15^) supposedly including various receptors. The distance to the transition state *x*_*u*_ provides a parameter to characterize the interacting cellular receptor as the distance between receptor and HA falls within this range. As an example, (**C**) shows snapshots of a force-probe molecular dynamics simulation between HA from influenza A/X31 (green) and a human-type receptor (red) (taken from ^15^). The distance between the terminal SA and Asn137 (magenta) is shown and scales between 4-20 Å. The measured values for *x*_*u*_ and *k*_*off*_ for the corresponding virus-cell combination are shown in **B** (green).

However, we observed contradicting preferences for H3N2/X31, AH1 and also for FPV. H3N2/X31 was found to preferentially bind avian-type cell surfaces, while only recognizing α-2,6-linked (human-type) receptors on the glycan array. AH1, similarly to pdmH1N1, recognized both receptor types on the glycan array and showed good binding to all six presented specific glycans on the array, while SVFS indicated a preference for human-type cell surfaces. FPV recognized all three avian, but only one human-type receptor on the glycan array, while showing preferential binding to human-type cell surfaces in SFVS. However, binding to the recognized human-type receptors (receptor 3 in Fig. 1) was about 2-3 fold stronger compared to the avian-type receptors (receptor 5 in Fig. 1), which might explain the stronger binding to A549 cells. In conclusion, our results suggest that the HAs receptor preference as tested in glycan array binding may not be a good predictor for preferred binding to human-type over avian-type cell surfaces.

The findings described above raised the possibility that non-sialic acid receptors contribute to a larger than expected extent to virus cell binding. However, SVFS analyses of cells after pre-treatment with neuraminidase to remove sialic acid structures showed that the binding probability was strongly reduced, leaving the unbinding force unchanged (Fig. 2B). This indicates that the viruses indeed mainly bind to sialic acid of the cell surface, but that the local environment of the receptor or other cell surface molecules alters the macroscopic cell specificity leading to the observed differences. The stronger binding of AH1, pdmH1N1 and H1N1/WSN to fucosylated glycan SLe^X^ with α-2,3-linked to SA in comparison to the other α-2,3-linked (avian-type) SA of our glycan array is indicative for the relevance of the local environment. On the viral side, cumulating evidence suggests a role of the viral neuraminidase (NA) in contributing to cell binding via sialic acid ^31^. In our SVFS measurements, NA was kept active and, hence, it cannot be excluded as a binding mediator, a feature that could be tested in future experiments.

We also took a closer look at the thermodynamic properties of the virus cell interaction (table 1). Comparison of the transition state distance *x*_*u*_ revealed values around 2-10 Å for viruses binding to A549 cells (mean 5.8 Å). In contrast, on CHO cells, we found higher transition state distances between 2-13 Å (mean 8.1 Å). This interesting feature was also previously observed for IAV H3N2/X-31^15^ and suggests a differently shaped energy barrier. Correlation of *x*_*u*_ and *k*_*off*_ revealed no apparent clustering of H1 or H7 viruses (Fig. 4A) indicating a dynamic interaction potentially involving multiple different receptor sites. However, some correlation can be observed suggesting that binding to human-type cell surfaces tends to result in lower bond energy and shorter unbinding distance. Indeed, *x*_*u*_ could be a possible parameter to explore structural differences underlying virus-cell specificity. The observed *x*_*u*_ values fall within the distance regime between receptor and its binding pocket. As an example, Fig. 4C shows snapshots of a force-distance molecular dynamics simulation between HA from influenza A H3N2/X31 and its human-type receptor (taken from ^15^). The distance between the terminal SA and Asn137 (magenta), part of the critical loop 130, is shown and scales between 4-20 Å. The corresponding value pair for *x*_*u*_ and *k*_*off*_ is shown in Fig. 3B (green).

## 4. Conclusion

Recent glycomics approaches and the use of *ex vivo* tissue culture revealed new insights into the complexity of the living cell surface ^13, 14^. Since sialic acid was first identified as an influenza virus attachment factor ^32^, many studies have focused on HA-SA binding. Although this interaction is clearly important, not only infection of desialylated cells ^11^, but also the recent characterization of non-SA binding hemagglutinin encoded by a bat-derived H17N10 virus^33^, and the discovery that 1918 pandemic virus unaffectedly binds to primary human airway cells even when its HA is engineered to bind exclusively to avian-type SA receptors ^34^ suggested that other molecular determinants within the plasma membrane are also critical in initiating influenza A virus infection. For characterizing virus specificity, we suggest a dual complementary approach: (1) *in vitro* binding assays with synthetic glycans to precisely identify the preference of HA (or NA) for a specific sialic acid structure and (2) SVFS as demonstrated here to unravel the cell specificity, modulated by the local environment of the living host cell. We have recently demonstrated this complementary approach for an adapted mutant of pdmH1N1 ^17^. While glycan array analysis could not identify a switch in receptor preference, SVFS revealed that the adaptive mutation in HA strongly reduced the binding strength without changing the cell specificity. These binding properties are not accessible and might be hidden when only using *in vitro* specificity assays. The use of new methods such as SVFS ^16^ and *ex vivo* tissue culture in combination with global glycomics and proteomics approaches could help to identify essential components of the plasma membrane facilitating influenza virus cell interaction.

## Materials and Methods

### Cell and virus propagation

Chinese hamster ovary (CHO) cells and human alveolar A549 cells were grown in DMEM (PAA) supplemented with 1% penicillin/streptomycin and 10% FCS (PAA) in plastic petri dishes. For sialic acid digestion, we used neuraminidase (NA) from *Clostridium perfringens* (Sigma) solved in PBS buffer. The cells were treated for 10 min at 37° C with 1 U/mL NA. Influenza A viruses were grown on 10-day old chicken eggs and purified from allantoic fluid by gradient centrifugation through a 20-60 % (w/v) sucrose gradient. The A/Anhui/1/2013 strain was inactivated by UV irradiation before gradient centrifugation.

### Glycan array

Glycan array preparation was performed as described previously ^35, Wormann, 2016 #192^. Briefly, glycans containing a primary amino linker were dissolved at a concentration of 0.1 mM in printing buffer (50 mM sodium phosphate, pH 8.5) and printed on N-hydroxysuccinimide activated glass slides (CodeLink slides, Surmodics, Edina, MN, USA) using an S3 robotic microarray spotter (Scienion, Berlin, Germany). Slides were incubated overnight in a humidity saturated chamber and remaining reactive groups were quenched by incubating with 100 mM ethanolamine, 50 mM sodium phosphate at pH 9.0 for 1 h at room temperature. Slides were washed with water, dried by centrifugation and stored at 4 °C until use. Before loading, the array was washed with DPBS. Virus was diluted as indicated into sterile binding buffer containing 1% BSA, 0.05% Tween 20 (MERCK), CaCl_2_ (492 μM) and MgCl_2_ (901 μM) at pH 7.0. 30 μl of diluted virus were pipetted in each well and the array was incubated in a moist chamber for 24 h at 4 °C. Each well was then washed three times with washing buffer containing DPBS and 0.1% Tween 20 (DPBS-T). Subsequently, wells were blocked with DPBS containing 1% BSA for 2 h at 4 °C and permeabilized using DPBS-T containing 0.3% Triton-X100. To stain the bound virus the array was incubated with a primary monoclonal antibody against the viral NP protein (1:1000, clone AA5H, AbD Serotec, Oxford, UK) at 4 °C overnight. Primary antibody was removed and wells were washed three times with DPBS-T. Secondary Cy3-coupled goat anti-mouse IgG (1:100, product-code: 115-165-146, Jackson ImmunoResearch Laboratories, West Grove, PA, USA) was added and incubated at RT for 1 h. The array was washed three times with DPBS-T and dipped into distilled water before scanning. Glycan array fluorescence images were obtained using a GenePix 4300A microarray scanner (Molecular Devices, Sunnyvale, CA, USA). Fluorescence intensities of spots were evaluated with GenePix Pro 7.2 (Molecular Devices).

### AFM tip chemistry

Commercially available AFM cantilevers (MSCT, Bruker) were amine functionalized by using the room-temperature method for reaction with APTES ^36^. A heterobifunctional PEG linker, acetal–PEG_800_–NHS (N-hydroxysuccinimide) (Fig. 1B), was attached by incubating the tip for 1.5-2 h in 0.5 mL of chloroform containing 2 mg/mL acetal-PEG-NHS and 8μL triethylamine, resulting in acylation of surface-linked APTES by the NHS group. The terminal acetal group was converted into an amine-reactive aldehyde by incubation in 1% citric acid as described previously ^36^. After rinsing with water for 3 times, once with ethanol and drying under a stream of nitrogen, the tips were incubated in a mixture of 19-25 μL of approximately 0.6-1.6 mg/mL influenza A virus in PBS (without Ca^++^) and 1-2 μL of 1 M NaCNBH_3_ (freshly prepared by dissolving 32 mg of solid NaCNBH_3_ in 500 μL of 10 mM NaOH) for 60 min. The tips were then washed in 3mL PBS for 3 times and stored in PBS at 4 °C. All other chemicals and reagents were purchased from different commercial sources in the highest purity grade available.

### SVFS measurement

As illustrated in Fig. 1, AFM-based force spectroscopy was performed with an Agilent 5500 AFM. The Petri dish with cells was mounted with the AFM, which was put on the optical microscope through a specially designed XY stage. Before force measurements, the cantilever with a nominal spring constant of 10 pN/m functionalized with influenza A virus was incubated in 5 mg/mL BSA for 30 min in order to minimize the nonspecific interaction between the cantilever tip and the cell surface. Measurements were performed in PBS buffer at room temperature. After the cantilever tip approached to the cell surface, force distance curves were repeatedly measured with Z-scanning range of 2 μm, cycle duration of 0.5-8 s, 500 data points per curve, and typical force limit of about 40-70 pN. The spring constants of the cantilevers were determined by using the thermal noise method ^37^.

### Fitting of SVFS data

Similar to single molecule force spectroscopy (SMFS), also in SVFS studies, several hundred force distance cycles are recorded in a dynamic range of increasing loading rates under identical conditions. For each of these force curves showing unbinding events, the unbinding force *Fi* and the effective spring constant *k*_*eff*_ (slope at rupture) were determined. The loading rates *r* were determined by multiplying the pulling velocity *v* with the effective spring constant *k*_*eff*_ (i.e. *r* = *v* * *k*_*eff*_). Additionally, a rupture force probability density function (pdf) (Fig. 1d) was calculated and a Gaussian distribution was fitted to the main peak of the pdf. Subsequently, all unbinding events within *μ* ± *σ* of the fit have been selected to create a loading rate dependence scatter plot (Fig. 1c-f) for further calculations of *k*_*off*_ and *x*_*u*_.

Generally, the loading rate *r* is constant for a fixed pulling speed, which implies, that the effective spring constant *k*_*eff*_ does not vary significantly. However, for force spectroscopy measurements on live cells it is known, that *k*_*eff*_ could show a broadened distribution caused by local variations of the spring constant of the cell surface, leading to a convolution of the rupture force distribution and further influences the calculations for the dissociation rate constant, *k*_*off*_, and the separation of the receptor-bound state to the energy barrier, *x*_*u*_. To circumvent this influence, we applied a maximum likelihood routine to fit the SVFS data to the Evans-model ^27^, in order to obtain *k*_*off*_ and *x*_*u*_ (Table 1).

Accordingly to the single energy barrier binding model, the probability *p* that the complex breaks at a certain force, *F*, is given as ^38^:

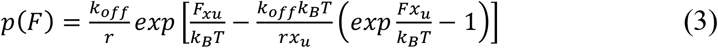

The parameters *x*_*u*_ and *k*_*off*_ were determined by applying a maximum likelihood approach, in which the negative log likelihood *nll* was minimized by modifying *k*_*off*_ and *x*_*u*_, with *p* based on Equation (3) defined in the single barrier model ^38^:

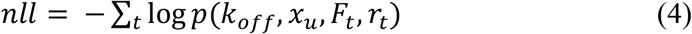

## Acknowledgments

This work was supported in part by the grants from the Hi-tech Research and Development Program of China (2008AA02Z311), the Shanghai Natural Science Foundation (13ZR1402400), the Shanghai Leading Academic Discipline Project (B111). The work was further supported by the German Ministry of Research and Education (BMBF) (e:Bio ViroSign) as well as the German Research Foundation (DFG) (SFB 765). R.Z. and P.H. were supported by Austrian Research Fund SFB-F35.

## Additional information

Supplementary information is available. Correspondence and requests for materials should be addressed to C.S., A.H. or P.H.

## Supplementary Figure Legends

**Supplementary Figure S1.**
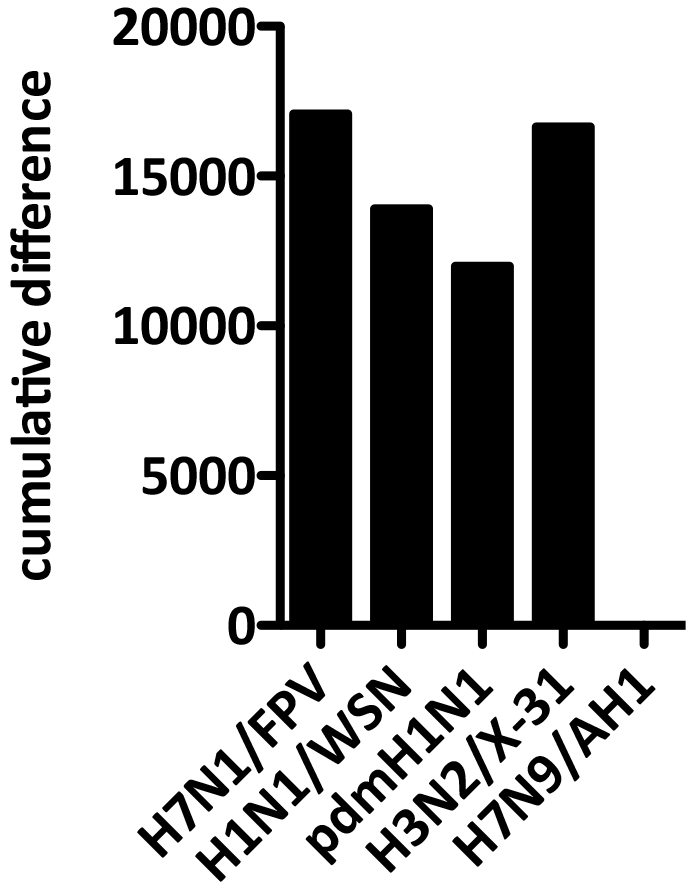
Cumulative receptor binding difference between H7N9/AH1 and the other on glycan arrays tested virus strains. For each glycan, the binding intensity was compared to the value of H7N9/AH1. The summed difference is shown for each virus strain revealing that H7N9/AH1 shares most similarities with pdmH1N1.

**Supplementary Figure S2.**
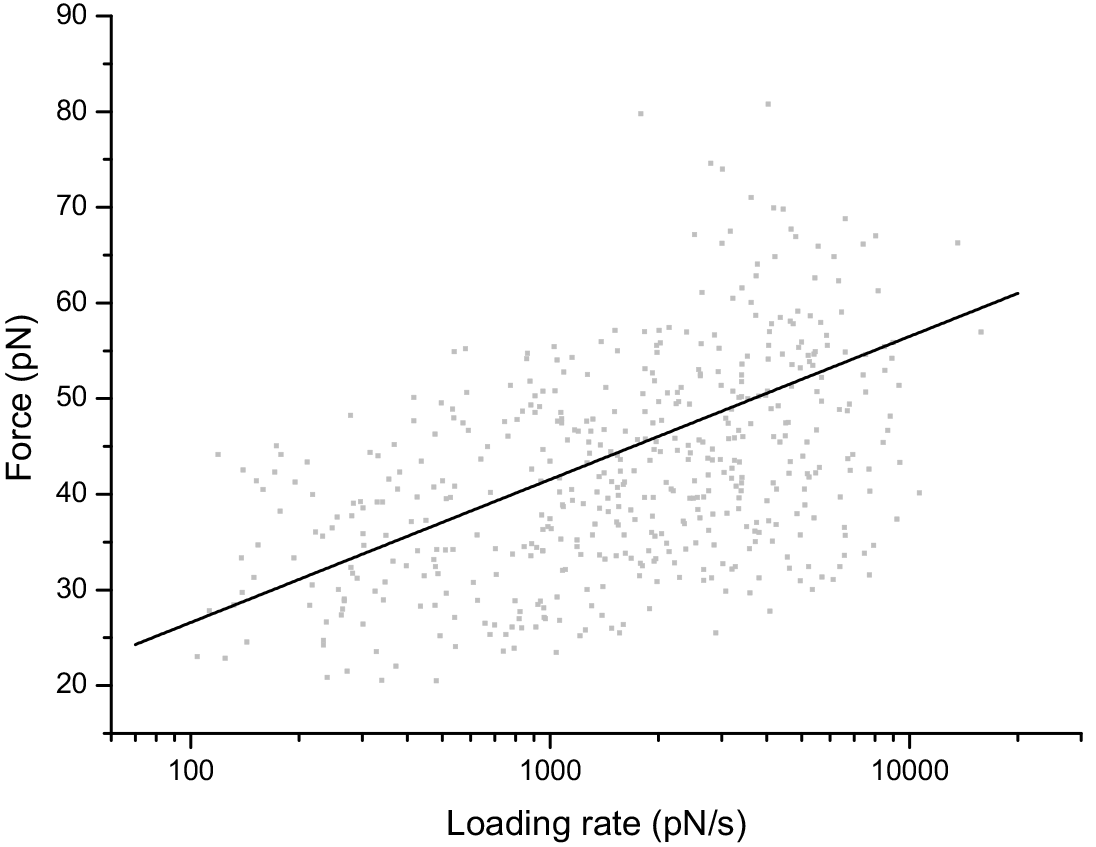
SVFS dynamic force spectra of H7N9 AH1 interacting with single receptors on living MDCK cells. Scatter plot showing unbinding force *F* plotted against the loading rate r of every individual force curve. From those data, the values for *k*_*off*_ and *x*_*u*_ were determined to be 0.256 +/− 0.00169 s^−1^ and 6.160 +/− 0.0146 Å, respectively.

**Supplementary Table S3.**
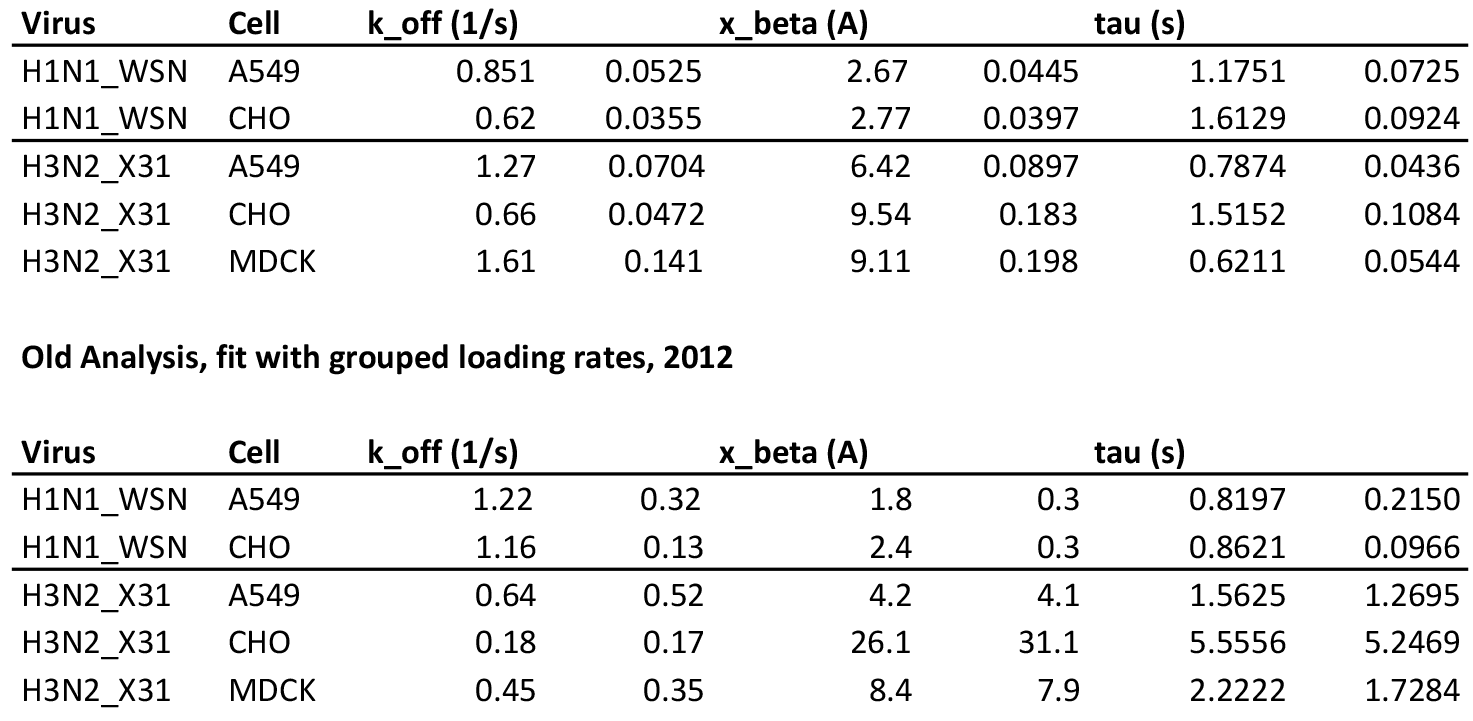
Comparing SVFS results obtained using two different fitting approaches. Loading rates *LR* can either be obtained by using the mean effective spring constant <*k*_*eff*_> for each pulling velocity *ν* (LR = v*k_eff_) or, using a more adapted approach reported previously and now used in this study, calculated for each individual force-distance curve. Each fitting approach results in slightly different fitting parameters. Dissociation rate *k*_off_, separation from the energy barrier *x*_u_, and average bond lifetime *τ*_off_ obtained by fitting the SVFS data to a single energy barrier binding model.

